# Analysis of Scholarly Discourse in Biosecurity from 1986-2020

**DOI:** 10.1101/2020.12.30.424860

**Authors:** Ye Henry Li, Milana Boukhman Trounce

## Abstract

Scholarly discourse in Biosecurity is complex and interdisciplinary. The term ‘Biosecurity’ has different meanings depending on the discipline. This term most often strives to capture efforts to prepare for and respond to threats posed by infectious organisms^1^. Increasing complexity of discourse in this interdisciplinary field and lack of common definitions can pose communication challenges. We use text mining of scholarly articles to characterize the interdisciplinary field of Biosecurity and the uses of this term over the past 35 years; a context evolution model is built to describe major changes in this field, and it shows a progression toward health security in recent years. Understanding the shifts in discourse predominance helps to illuminate the major challenges facing biosecurity research and academic community, and contributes to enhancing communication across disciplines within the Biosecurity field.

## Introduction

What is Biosecurity? The term has been defined differently by various disciplines, and the range of definitions is significant. For example, David Fidler and Larry Gostin, in their book “Biosecurity in the Global Age”, define biosecurity as “collective responsibility to safeguard the population from dangers presented by pathogenic microbes, whether intentionally released or naturally occurring’.^1^ World Health Organization defined it as “strategic and integrated approach to analysing and managing relevant risks to human, animal and plant life and health and associated risks for the environment”.^2^ One of the first uses of the term was in the context of agriculture^2^, where it referred to protecting animals and plants from risks posed by disease due to infectious organisms. It has also been used in the context of lab safety and biosafety, and involved issues such as setting regulatory guidelines surrounding the use of CRISPR technology^3^, mitigating the spread of the COVID-19^4^ and Ebola^5^, minimizing the loss of animal and plant species diversity from Australia’s devastating wildfires^6^, securing the smallpox virus at storage locations^7^, and many others. We examine academic discourse involving biosecurity through time and examine the evolution of this field and its major drivers.

Using text mining we propose evolutionary analysis of scholarly publications involving Biosecurity from 1986 through early 2020 to characterize this increasingly complex landscape. These findings give evidence that the Biosecurity field has progressed in different phases over the past 35 years. During this progression, while the new and old topics are intertwined through time, we observe a focus towards topics focusing on infectious diseases and a shift towards health security (both animal and human). It is likely that the ongoing COVID-19 pandemic will solidify Health Security as the dominant area of Biosecurity in the foreseeable future.

## Methodology

Google Scholar is a tool that enables searches of scholarly articles in scientific journals and books. Google Scholar returns the articles’ titles, year published, authors, publisher, the number of citations, and more. These search results are ranked by relevance based on a number of criteria, so it is possible to learn the general landscape of a field by surveying only the subset of top-ranked academic articles of that field without mining through every article ever published on the subject.

To data mine technical information efficiently, we use the citation tool Publish or Perish^8^ to collect all the top Google Scholar search results from 1986 through early 2020. Note that this search does not comprehensively include all of Biosecurity literature. We search for “Biosecurity” to extract the most Biosecurity-relevant articles. We divide the search cycles into 5-year periods starting 1986. Each search cycle yielded up to 1,000 top-ranked articles for each time period. In addition, to dissect “Dual-use” and “Health Security” (two dominant areas of study today), we search these terms in conjunction with “Biosecurity” to extract deeper insights. All searches and analyses were performed through May, 2020. The raw corpuses are available in the supplemental materials.

Downstream data analyses were performed using the R programming language^9^. From the search results, the publication titles were extracted and processed. In this work, we focused on the information gleaned from article titles with the presumption that they capture the main points of the articles succinctly. Note that analyzing corpuses from article abstracts and contents may yield additional layers of information to dissect areas of studies even further.

In the processing of article titles, we remove stop words, numbers, punctuations, and common words like *new* and *biosecurity* that occur in these searches that do not add more information to the content. The words are then condensed into word stems using the tm package in R.

To create snapshots of the Biosecurity landscape, the time axis from 1986 through 2020 is partitioned into seven 5-year time periods. We performed analysis on the top-ranked word stems across these periods, and built a model to describe the progression of Biosecurity discourse through time.

## Analysis and Discussion

The volume of Biosecurity scholarly articles has increased significantly over time (Figure 1). There are 494 raw results returned for the search of Biosecurity scholarly articles from the time period 1986-1990; consistently through subsequent periods, the number of relevant articles has increased drastically, ending with 27,400 in the current time period. We interpret this trend as that Biosecurity has gained significantly more intense academic focus over time. The highest percentage increase occurred in the 2001-2005 period, which corresponds to the scholarly mention of biosecurity along with issues concerning health, disease, and risk (Table 1). The numbers of relevant scholarly articles have been consistently higher in subsequent time periods. We have controlled for the number of articles by taking the 1,000 top-ranked search results from each period to extract key representative words for the comparison analysis.

**Table 1.**
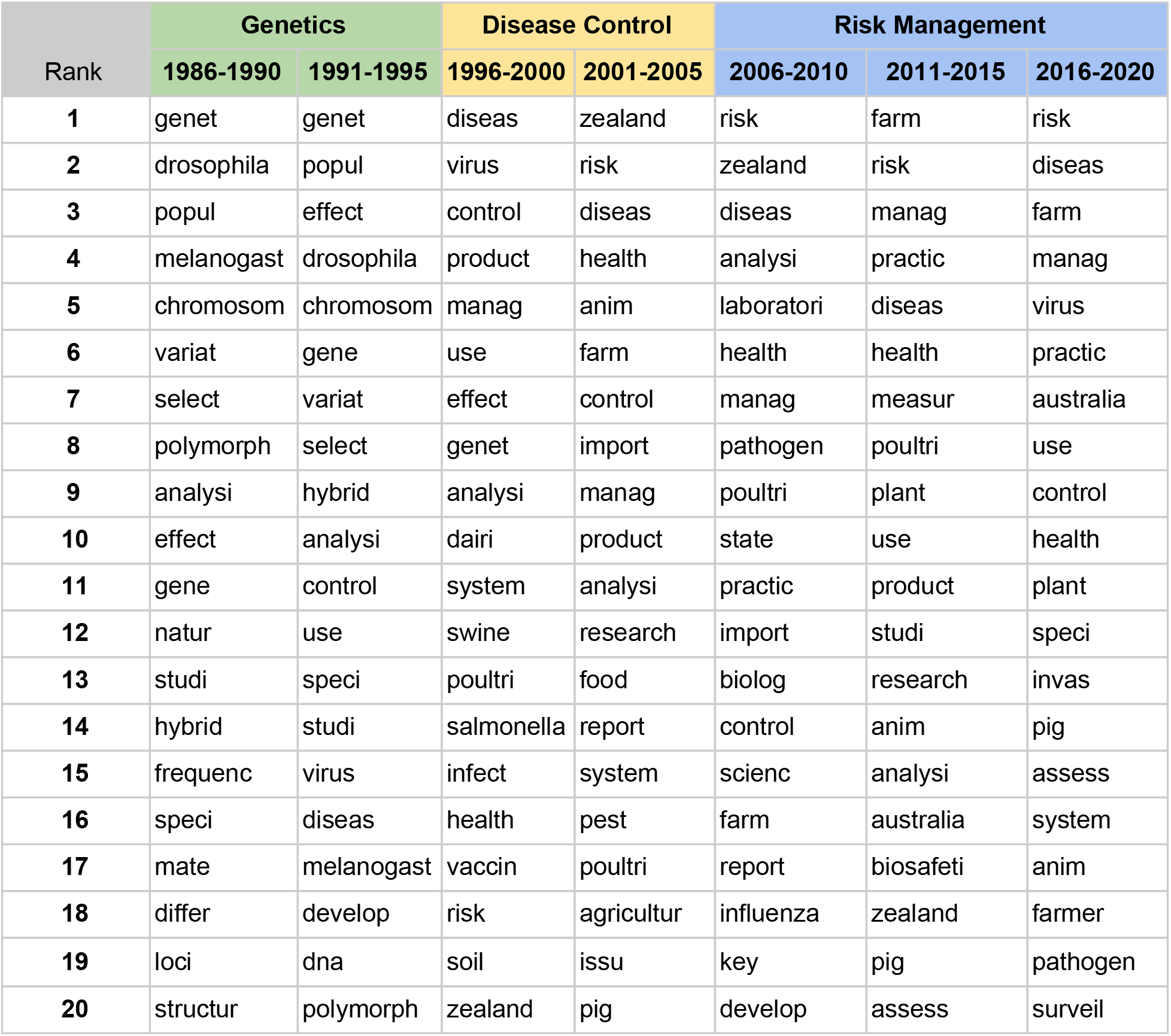
The top 20 ranked topic root words from each time period. As time progresses, new topics emerge as a result of the Biosecurity field becoming more diversified and complex. The periods are color-coded as three phases: green for population Genetics, yellow for Disease Control, and blue for Risk Management.

**Figure 1.**
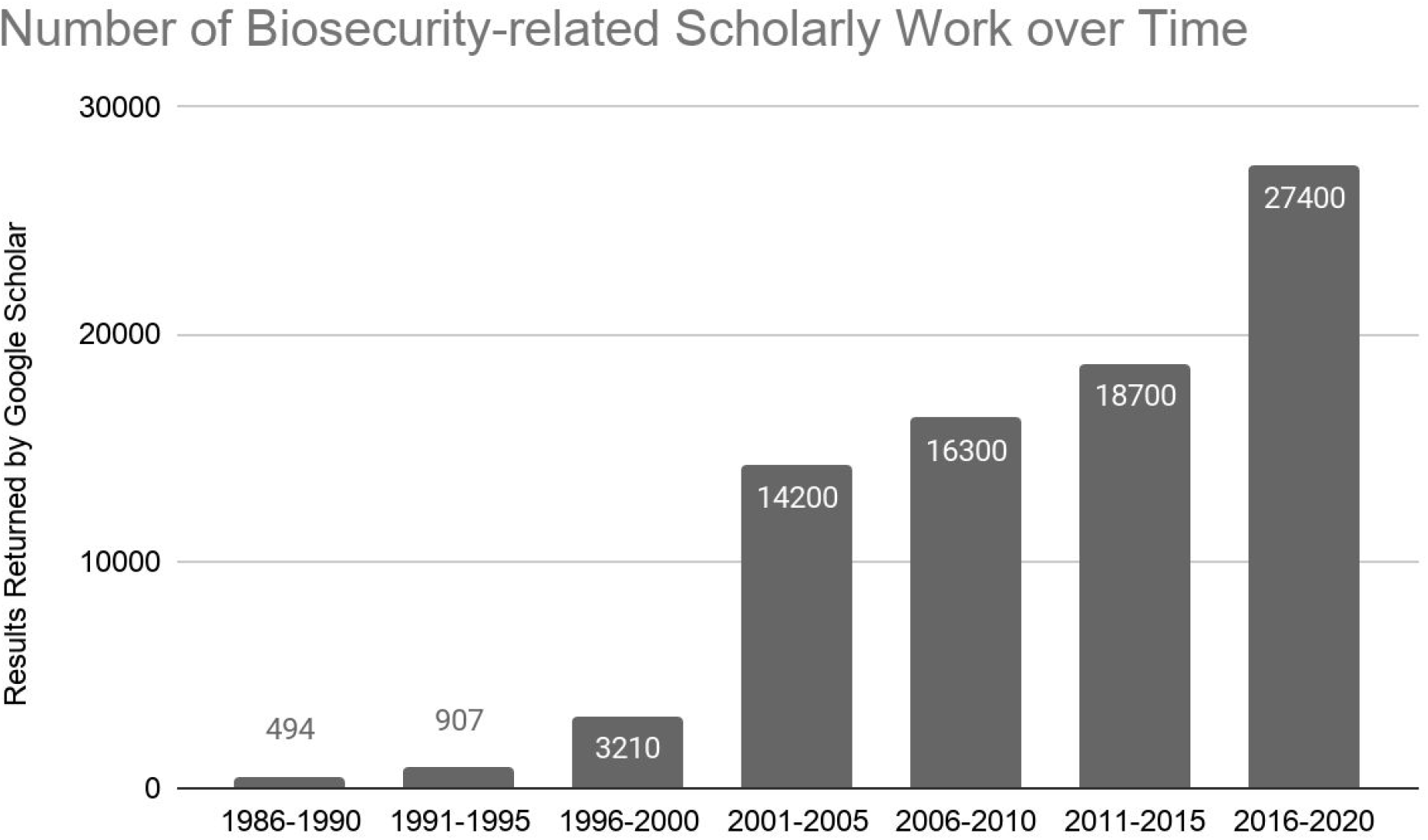
The numbers of raw articles returned in Google Scholar searches for 5-year time periods from 1986 through early 2020. There is a consistent increase over time, with the largest percentage jump occurring from 1996-2000 to 2001-2005.

From the analysis of the data involving top-ranked articles in these periods, we made two observations. First, academic discourse in Biosecurity changes through time. Second, the predominance of academic discourse has been intertwined as a result of the interdisciplinary nature of the field. We discuss these two observations in additional detail below. We also discuss the concerns surrounding dual-use technologies, one of the dominant areas of academic discourse in the field of biosecurity, and the emergence of Human Health Security as a major area of Biosecurity in recent years.

### Predominance of topics in phases

Through the last three decades, there have been increasing numbers of technical publications associated with Biosecurity, demonstrated by the increasing number of Google Scholar search results during each 5-year period (Table 1). Although Publish or Perish returns the top 1,000 Google Scholar search results, the titles of these technical publications can be used to construct corpuses for us to understand the overall Biosecurity topics through time.

The periods from 1986 through 1995 are dominated by Genetics. The top words from 1986-1995 are *genet, drosophila, popul, melanogast, chromosom, variat, select, polymorph, analysi*, and *effect*; and the top words from 1991-1995 are *genet, popul, effect, drosophila, chromosom, gene, variat, select, hybrid*, and *analysi*. These two five-year periods share high similarity in their top-ranked words. Most of Biosecurity-relevant works are on genetic diversities, such as studies on fly genetics and how plant alleles are inherited. Interestingly, an economic impact study of management practices to limit infections in chickens was conducted during this time^10^. This study on infectious diseases in chickens relates to the economic impacts of animal diseases on a much larger scale in the ensuing time periods^11^. These early periods are encapsulated by studies on animal and plant genetics as well as population genetics applications for agricultural sciences.

We observe an emergence of discourse centering on disease agents and control at the end of the last millenium. In the 1996-2000 period, a drastic increase has occurred in the number of scholarly articles related to Biosecurity, and these publications yield *diseas, virus, control, product, manag, use, effect, genet, analysi and dairi* as the set of high frequency stem words. Many articles in this period explore the relationship between agricultural practices and the threats of viruses, including avian flu viruses, that could derail the agriculture industry. For example, one article explains the set up of a pathogen-free system for shrimp aquaculture, in which there is water quality monitoring and a strategy to mitigate virus outbreaks^12^. Here, the focus of Biosecurity has shifted from Genetics to the investigations of disease agents, and the economic implications of agricultural failures.

At the start of the 21st century, we observe a coalescence of topics including risk management, environmental science, and agricultural practices. These intertwining topics are correlated with an explosion of nearly 500% increase (Figure 1) in Biosecurity-relevant articles compared to the preceding time period. In addition, around the years 1995-2005, the world has experienced an unprecedented and explosive growth in trade volume^13^, in which trade policies across the world encouraged raw materials and goods exchange to capture economic efficiencies.

Given these world events, it is reasonable that the top-ranked topics in the 2001-2005 time period are *zealand, risk, diseas, health, anim, farm, import, control, product*, and *manag*. One of the more Biosecurity-relevant articles from this period is about the development of DNA barcodes to provide rapid and accurate identification of morphologically indistinct invasive species; and this study envisions an application of the underlying technology for New Zealand to track species migration from the imports of goods^14^. Another top-ranked publication is on the strategies to protect New Zealand’s biome^15^. We interpret this shift in predominance of discourse as a result of more efficient applications of Genetics to agricultural sciences, which in turn has changed the shipments of goods around the world; with higher volume of movements across the continents, experts have looked more closely at the effects of the introduction of non-native species around the world. In addition, it is in this period that we observe the first mention of anthrax being tied to bioterrorism^16^, which corresponds to the act of bioterrorism the United States experienced^17^; bioterrorism topics will gain much more focus in the next period.

Finally, we entered the current phase of Risk Management. In the three time periods 2006-2010, 2011-2015, and 2016-early 2020, we observe similar sets of top words containing *risk, manag, disease*, and *health* consistently. In this phase, academic discourse has shifted toward dual-use technologies made in the laboratory, bioterrorism concerns, and the protection of native biomes from agricultural and trade practices. One top-ranked publication is a book on Biosecurity and Bioterrorism^18^. This signifies the collective concern by biosecurity experts that we have increasing anxiety for the regulation of our biological technologies in driving the global economy and our footprint in the world. For agricultural sciences, experts are taking a closer look into the use of antimicrobial treatments as it relates to biosecurity^19^. Others have analyzed control strategies in managing bacterial growth using feed additives and vaccination^20^. The academic discourse is reflective of the impact of globalization on changes in biomes around the world. Complicating risk management efforts is the arrival of increasingly sophisticated molecular technologies such as CRISPR genome editing tools. As such, the volume of scholarly activity in this phase has increased significantly.

### Intertwining topics through time

We can dissect the changes further as intertwining topics (Figure 2). Broadly, based on the top-ranked words from each period, there are several bundles, or categories, of topics: genetics, disease agent, agriculture, and risk management and control. They co-exist and co-evolve through time.

**Figure 2.**
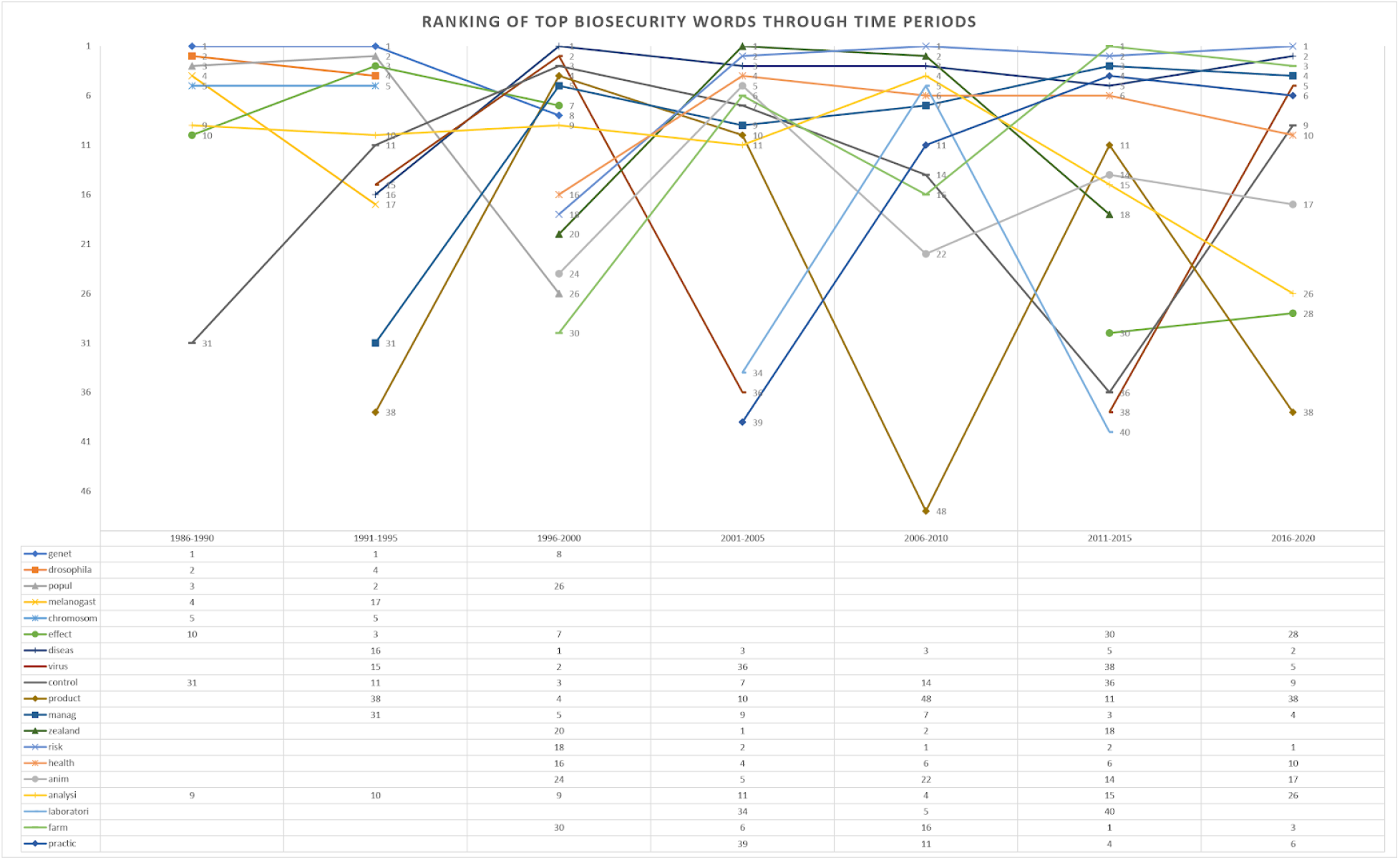
Top-ranked words shift in importance through time. While the predominance of topics has changed through time, major categories of academic discourse remain, and they intertwine.

The genetics category, which can be characterized by *genet, drosophila*, and *popul*, is more prominent in the years 1986-1995, though it has fizzled out by the start of the 21st century. This drastic decrease in importance may be due to the incorporation of a large volume of work from the disease control category as the Biosecurity-relevant literature increases by 300% in the 1996-2000 time period. Nevertheless, this category has set the foundation for the progression of subsequent predominant categories.

The disease agent category can be characterized by *diseas* and *virus*; it peaks in the Disease Control phase and continues strongly in the Risk Management phase. On the one hand, mentions of diseases and viruses in 1996 through 2005 come mostly from agricultural science articles, in which the major concern is about farm animal health. On the other hand, the mention of disease and viruses in the Risk Management phase, especially in the years 2016-2020, is from agents that affect human health. For example, there are articles about the biosecurity issues surrounding the smallpox virus^21^ and the presence of influenza virus (H5N1) in farm reservoirs^22^. Thus, while this category is characterized by disease agent issues, the underlying perspective has shifted from agriculture to human health over time.

The agricultural category can be characterized by *farm*, and it oscillates in its importance in relevance to Biosecurity as a whole. In the Disease Control phase, we observe the expansion of the focus from primarily animal health to human health, including articles that evaluate strategies and farming practices to protect farmers and the broader population^23^.

Risk management can be characterized by *risk, manag*, and *control*. This category is characterized by blending of elements from genetics, disease control, and agriculture categories. In combination with the other categories, it focuses on protecting animal and human health, with the ultimate goal of improving public health. The expansion of risk management coincides with an increased importance of *health* during these time periods. In addition, we note that many topics coexist in the Risk Management phase, suggesting that experts are reading and discussing a wide-ranging spectrum of Biosecurity issues.

### Dual-use technology concerns

In the 1986-1990 period, we observe Genetics is the predominant area of study in Biosecurity. Just a decade prior, in 1975, the ground-breaking Asilomar Conference set up guidelines for scientists to pursue recombinant DNA research while protecting public safety^24^. Recently, with the advent of more efficient DNA modification technologies such as CRISPR-Cas9, dual-use technologies have dominated academic discourse^25^. We have found *research, biolog, scienc, secur, life, ethic, synthet, biosafeti, concern*, and *risk* as the top 10 words for the dual-use area in the 2011-2020 period (Figure 3). Issues of ethics and security in the biological sciences dominate the academic discourse in regards to issues of dual-use.

**Figure 3.**
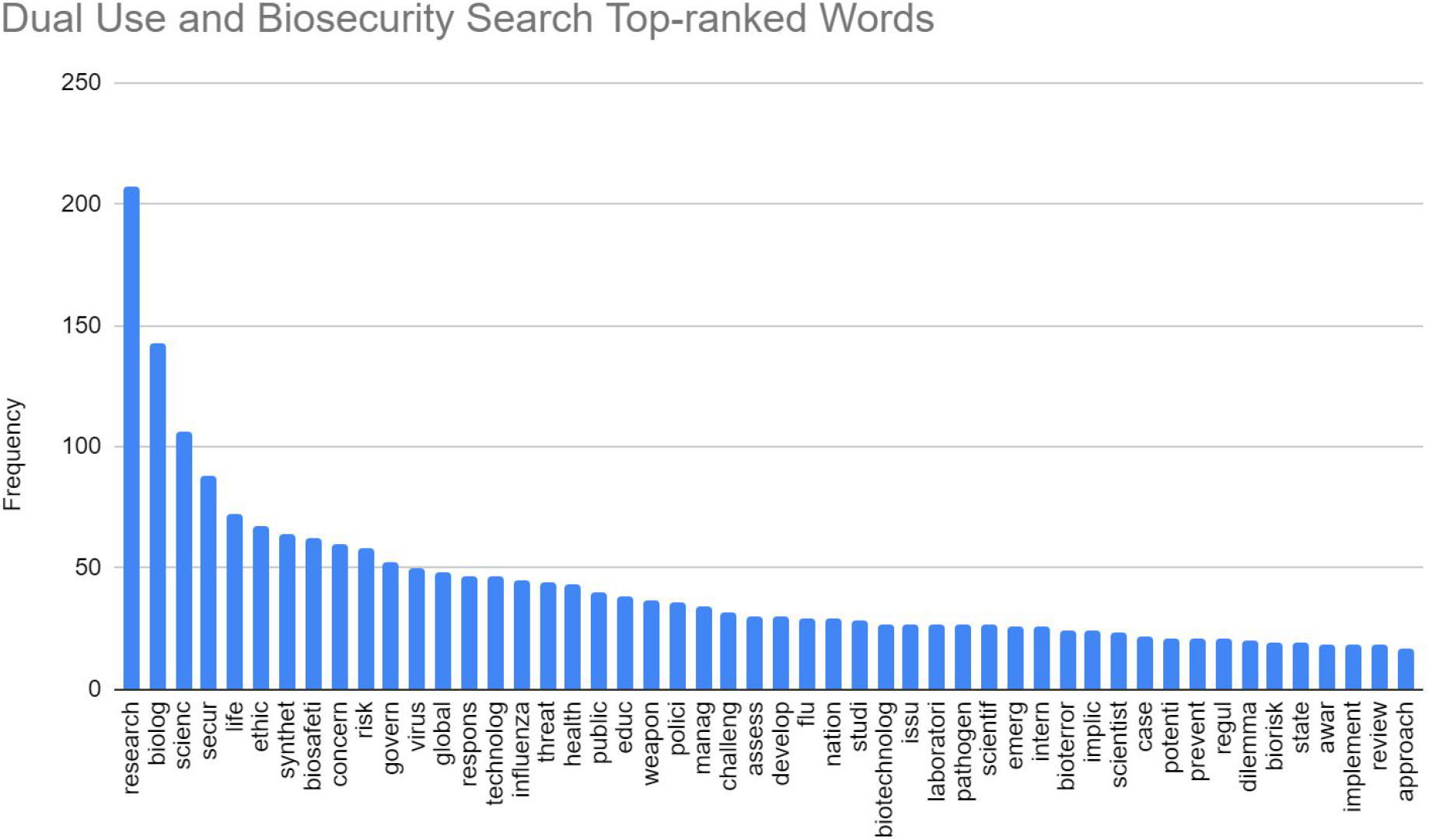
The top-ranked root words from scholar search using *dual use* and *biosecurity* for the period 2011-2020.

While dual-use discussions have risen in prominence, there are numerous other areas within BioSecurity that are strongly related to risk management.

### Human health security

As mentioned earlier, the amount of academic discourse focused on health security, as it relates to both animal and human public health, has increased in recent years. We examine the discourse in Health Security to understand its relationship with the changing Biosecurity landscape.

From the top-ranked word analysis, *global, diseas, biosafeti, public, manag, biolog, research, laboratori, risk*, and *food* are found to be the most dominant (Figure 4) in Health Security. In addition, the term *global* occurs frequently in scholarly articles, almost three times more often than the next top word *diseas*. The importance of *risk* and *manag* here aligns with the context of Risk Management. We may be looking at a shift of Biosecurity into one that is dominated by Health Security.

**Figure 4.**
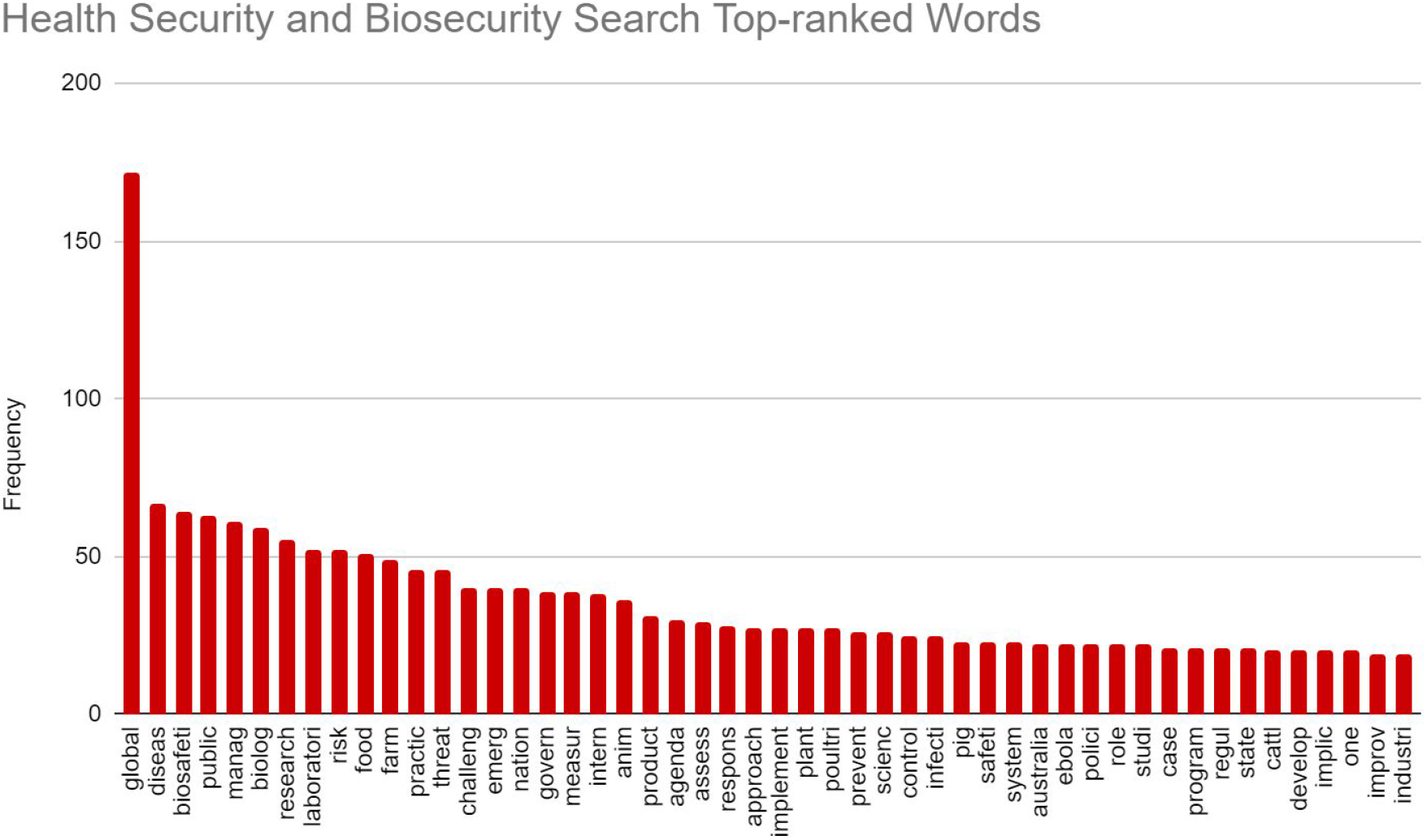
The top-ranked root words from scholar search using *health security* and *biosecurity* for the period 2011-2020.

The World Health Organization is addressing Health Security issues as part of global preparedness for potential deadly disease outbreaks^26^. The ongoing COVID-19 pandemic as well as other infectious disease outbreaks such as Ebola, SARS, MERS and others have propelled Health Security to be the dominant area of a more recent discourse.

## Conclusion

Using a text mining approach with context evolution analysis, we have quantified aspects of the use of the term ‘Biosecurity’ in academic literature and illustrated the evolution of academic discourse relevant to Biosecurity over time. We show that progression of the Biosecurity landscape is dynamic and complex, and its scholarly focus has changed through predominant phases that contain branches of intertwining topics as outlined above. We have shown that academic discourse in Biosecurity expanded many-fold over the past three decades and illustrated the emergence and increasing dominance of new areas of focus such as dual-use and human health security. We hope that this analysis provides perspective on the evolution of the field of Biosecurity and the uses of this term. We also hope that this understanding will contribute to enhancing interdisciplinary communication in this field and thus facilitate the creation of solutions to the many challenges posed by the threat of infectious organisms.

